# Synergistic effects of tRNA modification defects in *Escherichia coli* K12

**DOI:** 10.1101/2024.11.12.622971

**Authors:** Jo Marie Bacusmo, Jill Babor, Jennifer Hu, Bo Cao, Stefanie Kaiser, Sierra Szkrybalo, Yifeng Yuan, Serena Sander, Paul Kuipers, Laura Sanchez Baldoquin, Michael DeMott, Hirotada Mori, Peter Dedon, Valérie de Crécy-Lagard

**Affiliations:** Microbiology and Cell Science, University of Florida, Gainesville, FL 32611, USA; Department of Biological Engineering, Massachusetts Institute of Technology, Cambridge, MA 02139, USA; Innovation Laboratory of Systems Microbiology and Synthetic Biology, Institute of Animal Sciences, Guangdong Academy of Agricultural Sciences, Guangzhou, Guangdong 510640, China; Antimicrobial Resistance Interdisciplinary Research Group, Singapore-MIT Alliance for Research and Technology, Singapore, Singapore; Genetics Institute, University of Florida, Gainesville, FL 32611, USA

**Author notes:** Molecular Biology Program, Sloan-Kettering Institute, New York, NY 10065, USA. CG Onocology, Irvine, CA 92618, USA. Goethe University Frankfurt, Faculty 14, Institute of Pharmaceutical Chemistry, Frankfurt 60438, Germany (Stefanie Kaiser). Shanghai Jiao Tong University School of Life Sciences and Biotechnology, 800 Dongchuan Road, Minhang District, Shanghai 200240, China (Bo Cao).

**Keywords:** tRNA modification, tRNA function, synthetic lethality, s^2^U

## Abstract

Transfer RNAs (tRNAs) are essential components of the translation machinery and carry numerous post-transcriptional modifications that contribute to decoding accuracy, efficiency and cellular fitness. In *Escherichia coli* K-12, all tRNA modification pathways have been identified, yet the functional interactions between these pathways remain largely unexplored. Here, we systematically analyze genetic interactions between 29 non-essential tRNA modification genes using a pairwise synthetic lethal screen based on P1 transduction. Most combinations of tRNA modification gene deletions are tolerated during growth in rich medium; however, we identify five synthetically lethal pairs and fifteen additional combinations that display negative genetic interactions. Deletions of *truA*, which encodes the pseudouridine synthase responsible for modifications at positions 38–40 of multiple tRNAs, show the highest frequency of negative epistasis. Synthetic lethality associated with *truA* can be complemented by expression of *truA* in trans and, in specific cases, partially suppressed by overexpression of its tRNA substrates, indicating substrate-specific functional dependencies. Analysis of tRNA abundance by northern blotting and AQRNA-seq demonstrates that loss of individual tRNA modification enzymes does not generally lead to widespread tRNA destabilization. Instead, further phenotypic characterization of viable double mutants reveals condition-dependent growth defects influenced by carbon source, temperature and metabolic stress, as well as toxicity associated with overexpression of specific tRNAs. Together, these results reveal a limited but distinct set of genetic interactions among bacterial tRNA modification pathways and highlight the importance of physiological context in uncovering their cellular roles.

## Introduction

Translation is an extremely intricate process with several key components playing integral roles in maintaining the fidelity and efficiency of the decoding machinery. At the core of this operation are tRNAs, which act as adaptor molecules bridging the nucleic acid and amino acid codes. tRNAs also participate in biological processes beyond translation such as reverse-transcription primer for some retroviruses, and amino acid donors in the biosynthesis of peptidoglycan, antimicrobial molecules, and membrane lipids [1–3]. To fulfill their multiple cellular functions, tRNAs are extensively modified [4]. These modifications display a wide variety of chemical complexity ranging from basic methylations to elaborate modifications like queuosine or wyosine [5]. The functions of tRNA modification are varied. They allow tRNAs to decode mRNAs accurately and efficiently [5–7], stabilize their tertiary structure [8], and act as determinants for other components of the translation apparatus such as aminoacyl-tRNA synthetase [8] or ribonuclease [9] to name a few. Further, tRNA modification profiles have been shown to respond to stress and changes in physiological conditions of the cell [10]. This adaptive response allows for the translational regulation of gene expression pertinent to specific stresses or cellular demands [11,12]. It has been well established for over twenty years that in yeast RNA modifications serve as checkpoints for RNA integrity as tRNA lacking one (or more) modifications are degraded by different tRNA surveillance pathways [12–15]. More recently, it was shown that the absence of tRNA modifications could also lead to degradation of bacterial tRNAs. In *Vibrio cholerae*, tRNAs lacking s^4^U are rapidly degraded in stationary phase in degradosome-dependent manner [16]. In *Salmonella typhimurium* LT2, the absence of Ψ38-40 in tRNA can lead to its cleavage and to the activation of a repair pathway [17].

Synthetic lethality screens have been effective in dissecting the role of tRNA modifications in tRNA quality control pathways. The first example was in yeast where a combination of 10 pairs of tRNA modification mutants were analyzed and led to the discovery of quality control tRNA degradation pathways [15,18]. The second was in *V. cholerae* where a TnSeq analysis of a *thiI* strain deficient in s^4^U led to the identification of negative genetic interaction with the *truA*, *miaA* and *truA* that revealed that the degradosome targeted unmodified tRNAs [16]. However, no synthetic lethal screens have, to date, focused solely on a whole set of bacterial tRNA modification genes. *Escherichia coli* is the most studied bacterial model gram-negative and numerous synthetic lethal screens have been conducted in this organism. Some screens focus on specific groups of genes such as transcription factors [19] or small RNAs [20]. Other screens focused on functional areas such as DNA repair [21] or translation [22]. Only 14 modification genes out of the 43 known in *E. coli* [23] were included in the latter study but no synthetic lethal pairing of tRNA modification genes were identified in this data set. To palliate the absence of systematic study, we therefore conducted a synthetic lethal screen of 29 tRNA modification genes in the model *E. coli*.

## Results

### tRNA modification profiles of 29 tRNA modification mutants confirm expected phenotypes but does not reveal major cross-talks

All *E. coli* modification genes have now been identified [24] but a few were still missing when we performed this study, thus were excluded in our analysis (underlined in **Fig. 1A**). Further, several genes are essential (in green in **Fig. 1A**) or involved in the same pathway such as queuosine. We chose a set of 31 genes covering 26 nucleoside modifications at 20 positions along the tRNA (in red in **Fig. 1A**) and obtained the corresponding mutants from the Keio collection (**Table S1**). Bulk tRNA was extracted and purified from 3 biological replicates of each of the *E. coli* BW25113 derivatives devoid of the 31 specific tRNA modifying genes. The tRNA modification profiles were then analyzed via mass spectrometry. As expected, we saw a significant decrease in the levels of a given modification when the gene responsible for its synthesis was deleted (**Fig. 1B, Supplementary data S1)**: ac^4^C was absent in Δ*tmcA*, (m)cmo^5^U in Δ*cmoAB*, i^6^A in Δ*miaA*,, m^2^A in Δ*rlmN,* m^6^A in Δ*trmM*, Gm in Δ*trmH*, m^5^U in Δ*trmA*, s^2^C in Δ*ttcA*, s^4^U in Δ*thiI*, and m^6^t^6^A in Δ*trmO*. A significant decrease in both epoxy-Q (oQ) and Q was observed in Δ*tgt* because Tgt is responsible for the transfer of preQ_1_ (precursor to both oQ and Q).

**Figure 1.**
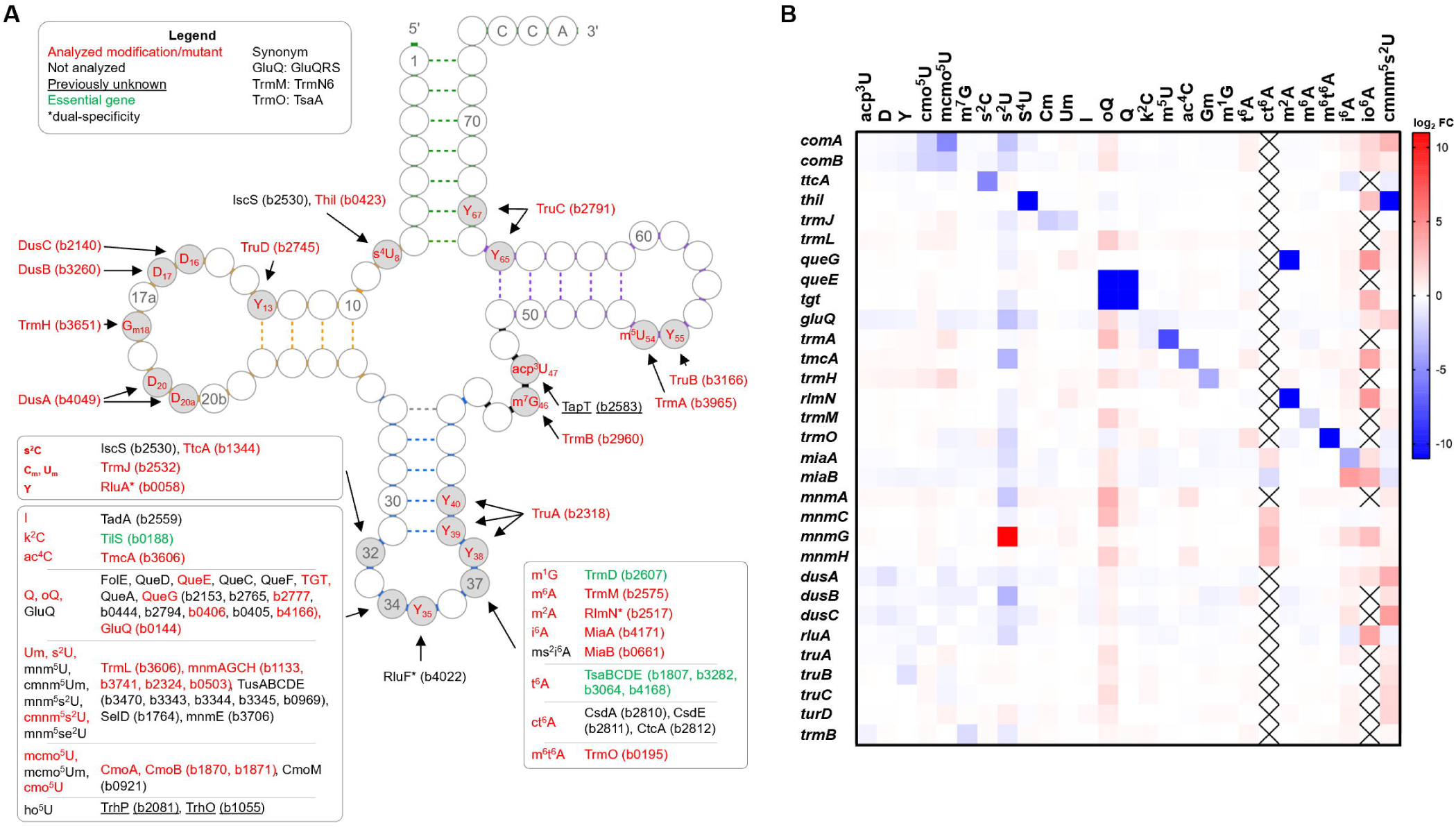
tRNA modifications proteins in *E. coli* K12 strain MG1655 and LC-MS validation of mutants analyzed in this study. (**A**) Positions of tRNA modifications and associated proteins and genes (blattner names are shown in parentheses). Modifications that were targeted are in red. Those inserted by essential genes are in green. Genes that were not identified at the time of the study are underlined. *: also modifies rRNA. (**B**) tRNA modification profiles in 31 different tRNA modification deficient strains. The genotypes of the Keio collection tRNA modification mutant strain are listed in **Table S1**. Modification abbreviations are based on MODOMICS (http://genesilico.pl/modomics/modifications). Data are presented as log2 fold change values relative to wild-type *E. coli* subjected to hierarchical clustering analysis. “X” denotes analyses in which no detectable signals were obtained for the modification. io^6^A that does not exist in *E. coli,* it was analyzed as a control.

We also saw expected trends in cases where several genes are responsible for one modification, as in the case of dihydrouridine (D) and Cm/Um. Most of the D modifications in tRNA are contributed by *dusA* and *dusB* while a smaller subset was modified by *dusC* [25]. Our results reflect this as we observe a 5-fold decrease in modification levels when *dusA* and *dusB* are deleted compared to when only *dusC* is deleted (**Supplementary data S1**). Modification of 2’-O-methylated cytidine (C_m_) or 2’-O-methylated uridine (U_m_) at position 32 in tRNA is catalyzed by *trmJ* (to tRNA^fMet^_CAU_, tRNA^Trp^_CCA_, tRNA^Gln^, tRNA^Gln^, tRNA^Ser^_UGA_) [26] and *trmL* (to tRNA^Leu^_CAA_ and tRNA^Leu^_UAA_) [27]. Our results reflect difference in the target tRNA populations of the two genes, with approximately a 16-fold decrease in C_m_/U_m_ modification levels observed upon deletion of *trmJ* compared with deletion of *trmL* (**Supplementary data S1**).

We also observed precursor accumulation upon deletion of genes responsible for downstream reactions in the pathway, as seen for Δ*miaB*, Δ*queG*, and Δ*mnmG* (**Fig.1B**). MiaB catalyzes the reaction that converts i^6^A into ms^2^i^6^A [28], thus an accumulation of i^6^A is observed in its absence. Similarly, an accumulation of oQ is observed in the Δ*queG* strain, as the missing enzyme catalyzes the conversion of oQ to Q [29]. Finally, an accumulation of s^2^U was observed in the Δ*mnmG* strain as the MnmG*-*MnmE complex converts s^2^U into either cmnm^5^s^2^U or nm^5^s^2^U [30]. Altogether, these observations validate our data set on each of the tRNA modification profiles.

Previous studies reported that i^6^A37 or ms^2^i^6^A37 stimulate U_m_34 and C_m_34 formation in *E. coli* tRNA^Leu^_UAA_ and tRNA^Leu^_CAA_ [31]. We did not observe any reduction in C_m_ levels in the *miaA* mutant but as we stated above, this could be because TrmJ-dependent methylations are still present in this strain.

One unexpected result was the disappearance of cmnm^5^s^2^U in the *thiI* mutant (**Fig. 1B, Supplementary data S1**). The cmnm^5^s^2^U modification is found in tRNA^Gln^_UUG_ at position 34. Its synthesis requires three enzymes: MnmEG for the formation of cmnm^5^U and MnmA for the thiolation step [32]. ThiI is the enzyme involved in the synthesis of s^4^U at position 8 found in many tRNAs including tRNA^Gln^_UUG_ [33]. The cysteine desulfurase, IscS, provides the sulfur moiety to both ThiI and MnmA [34]. tRNA^Lys^_UUU_ does not contain s^4^U while tRNA^Gln^_UUG_ does [33]. No obvious explanation for this observation could be proposed. If there is competition for the sulfur source, then one would expect that the absence of ThiI would lead to increased and not decreased s^2^U levels. If s^4^U is determinant for MnmA then only tRNA^Gln^ should be affected and there should be a reduction not an elimination of cmnm^5^s^2^U.

### Evaluation of epistatic relationships between tRNA modifying genes reveal a subset of genes with higher propensity towards synthetic lethality

To explore epistatic relationships between tRNA modifying genes, a synthetic lethal screen was conducted. We generated a pairwise deletion matrix of 29 tRNA modifying genes covering 26 different modifications, excluding *cmoA* and *queE* from the previous set of 31 genes, via P1 transduction. The matrix was evaluated based on the number of transductants obtained on LB plates at 37°C relative to the wild-type control. Two sets of deletion strains were used as donors and recipients for the P1 transduction: one set contained chloramphenicol resistance cassette (cat) inserted in the place of the deleted tRNA modification gene, while the second set contained a kanamycin resistance cassette (kan, strains were taken from the Keio collection). The list of the strains used is provided in **Table S1**. Transductions were performed in either direction selecting for transductants resistant to both chloramphenicol and kanamycin. All mutant combinations exhibiting no transductants, less transductants, or transductants with smaller colony sizes were repeated independently (**Supplementary Data S2**). We also verified that none of the gene pairs showing growth-defect phenotypes were within co-transduction distance, which could otherwise lead to artifacts (**Fig. S1**). The synthetic lethal screen showed that 95% (386 of 406 pairs) of the tested combination of deletions of tRNA modification genes gave WT looking colonies, while 4% (15 pairs) showed synthetic defects giving fewer transductants or transductants with smaller colony sizes and 1% (5 pairs) resulted in synthetic lethality (**Fig. 2**). Seven genes appeared in synthetic lethal pairs: Δ*truA,* Δ*mnmA,* Δ*rlmN,* Δ*rluA,* Δ*miaA,* Δ*truD*, and Δ*trmJ.* Out of the five synthetic lethal pairs, four encode two enzymes modifying the Anticodon Stem Loop (ASL) (Δ*rlmN*Δ*miaA*, Δ*truA*Δ*mnmA*, Δ*truA*Δ*rlmN*, and Δ*trmJ*Δ*truA*) while one pair encodes enzymes modifying the D-arm and the ASL respectively (Δ*truD*Δ*rluA*) (**Fig. S2**). The MnmA/MnmG synthetic lethality had been previously reported in *S. enterica* LT2 [35] and was reproduced here in *E. coli*.

**Figure 2.**
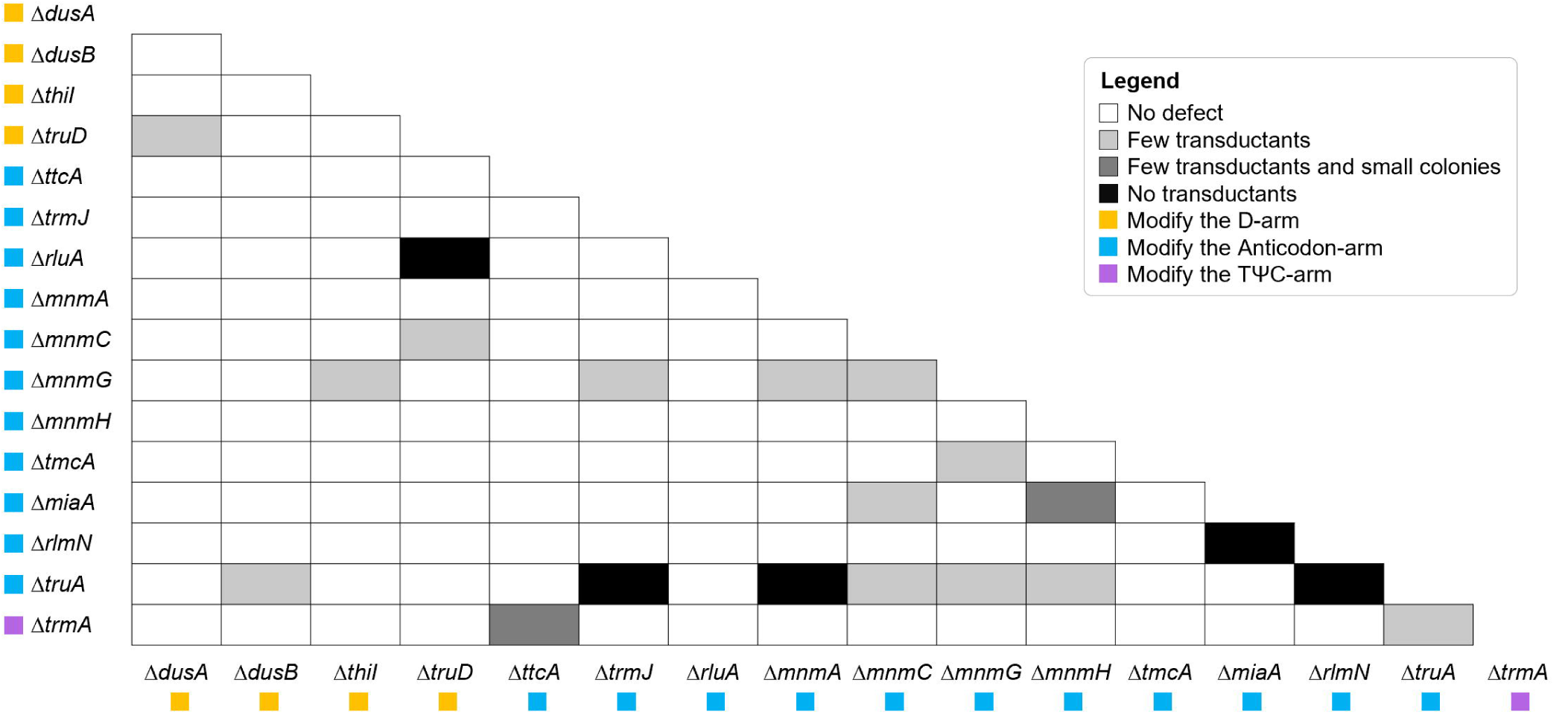
Synthetic lethality screen of 16 tRNA modification genes. All the mutant combinations that intersect with white boxes gave rise to transductant numbers equivalent to transducing the WT strain. The combinations that gave rise to growth phenotypes are noted by blank (no defect), light grey (fewer transductants), medium grey (small colonies and fewer transductants), and black (no transductant) boxes. The positions of modifications introduced by each gene are indicated by yellow (D-arm), blue (anticodon-arm) and purple boxes (TΨC-arm), respectively. The mutant combinations exhibiting growth-defect phenotypes on an initial screen were confirmed individually (**Supplementary data S2**).

The highest percentage of negative epistasis was observed when transducing any of the *dusB*, *mnmA*, *mnmC*, *mnmG*, *mnmH*, *rlmN*, *trmA* or *trmJ* deletion alleles into the Δ*truA* strain with a variety of phenotypes (small colonies, fewer or no colonies). As all the attempts to obtain Δ*truA*Δ*mnmA* clones failed, we studied this synthetic lethality phenotype further by measuring the P1 transduction efficiency of the Δ*mnmA*::*cat* allele into *truA* deficient derivative recipients. Transductants were only obtained when *truA* was expressed *in trans* and adding inducer led to more transductants. (**Fig. 3A**). Similarly, expressing the *truA* gene i*n trans* complemented the epistatic growth defects fully (for *dusB*, *mnmH* and *trmJ*) or partially (for *mnmC, trmA* and *rlmN*) (**Fig. S3**). The only tRNA that requires both MnmA and TruA to be fully modified is tRNA^Gln1^_UUG_ (**Fig. 4**) and indeed, expressing this tRNA (and not tRNA^Gln2^_CUG_) in trans partially suppressed the Δ*truA* Δ*mnmA* synthetic lethality (**Fig. 3B**).

**Figure 3.**
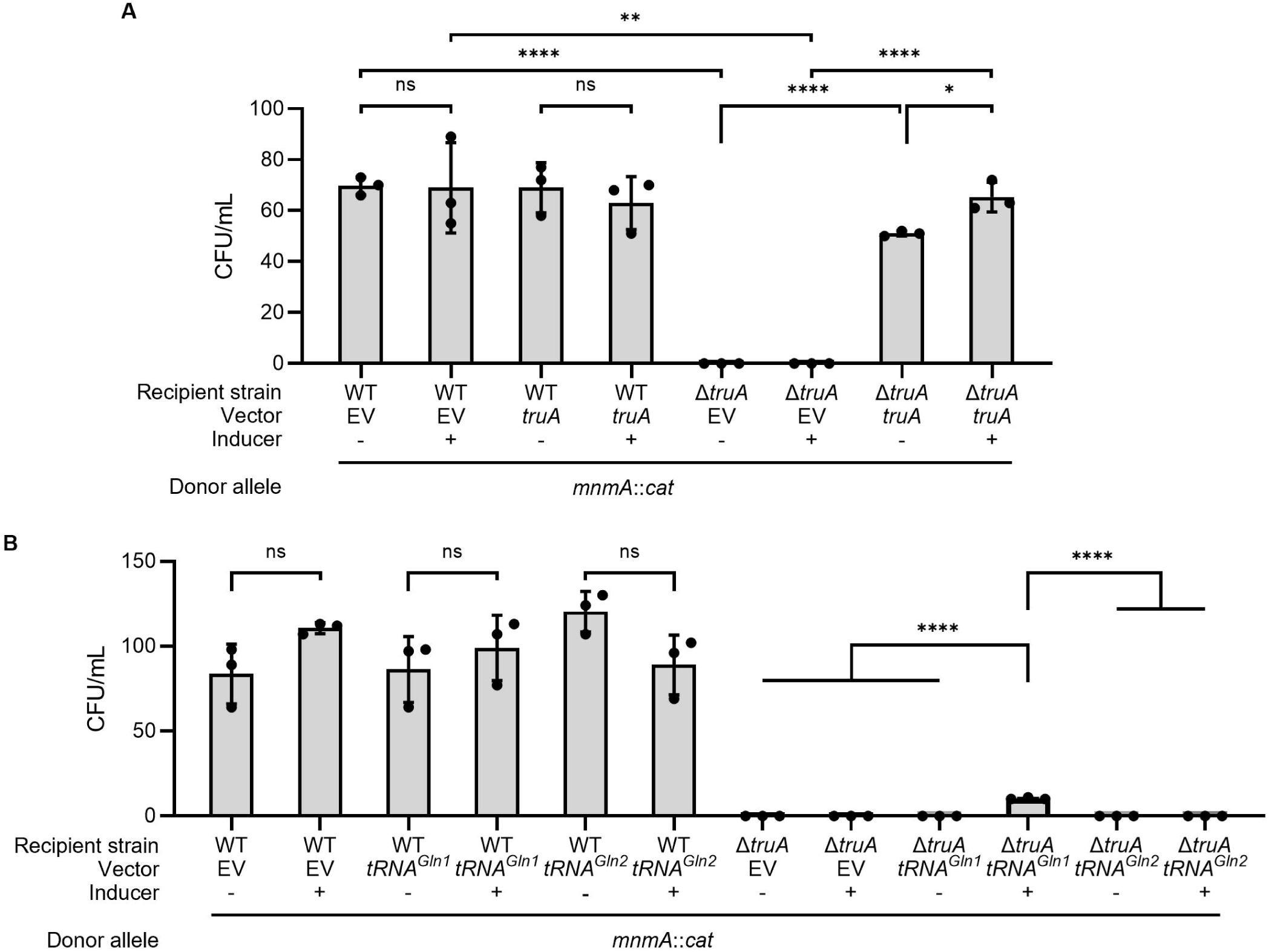
The synthetic lethality phenotype of the *truA* and *mnmA* genes is complemented by expressing the *truA* gene in trans and partially suppressed by overexpressing tRNA^Gln1^. P1 transduction efficiency was measured by the number of transductants when selecting for the insertion of the Δ*mnmA*::*cat* allele into the WT or Δ*truA*::*kan* strains after 24h incubation at 37°C. The synthetic lethality phenotype is complemented by a pBAD plasmid expressing *truA* gene (**A**) and a pTrc99a plasmid expressing *tRNA^Gln1^* (**B**). The selection for transductants was performed on LB plates supplemented with ampicillin (200 µg/mL), and chloramphenicol (25 µg/mL) with arabinose (0.02%) or IPTG (0.1 mM) as inducer for pBAD and pTrc99a plasmids respectively. An empty vector (EV) and no inducer was used as control. The average numbers of three replicates were plotted with error bars representing standard deviation. ns: P value < 0.05, *: P value ≤ 0.05, **: P value ≤ 0.001, ****: P value ≤ 0.0001.

**Figure 4.**
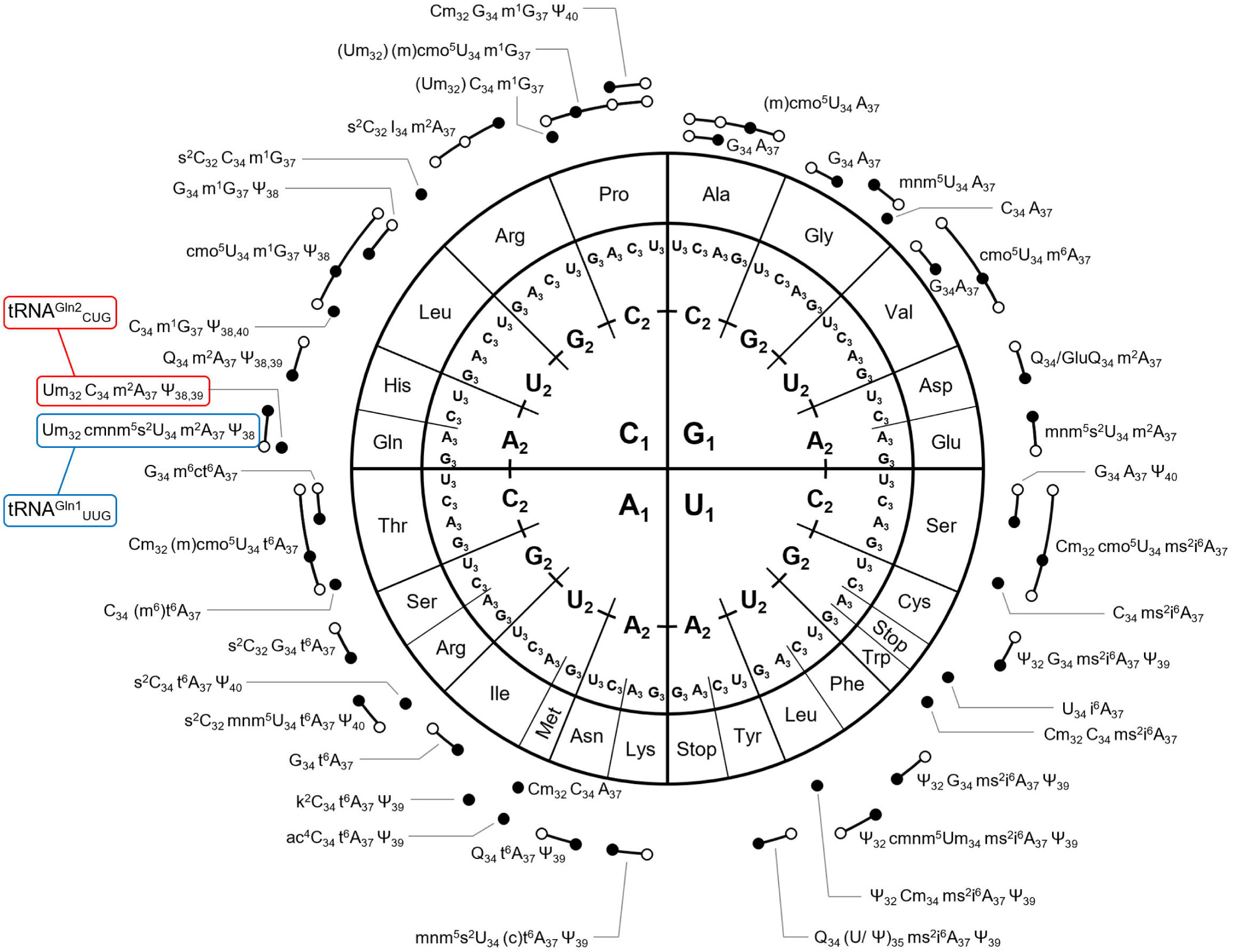
The tRNA decoding capacities and ASL modifications of *E. coli*. Letters from inside out indicate the codon’s first, second, and third position (5′- to 3′-end) as the subscripts indicate. The circles outside of the wheel connected or not by a line represent one tRNA species. A filled circle indicates the capacity of that tRNA to base pair with the indicated codon by Watson-Crick pairing. An open (white) circle suggests base pairing by the wobble hypothesis [50]. The base at position 34 of the anticodon and the anticodon stem loop (ASL) modifications in positions 32 to 40 are shown next to each tRNA circle. Amino acids that are decoded by a single tRNA are highlighted in red. Data are compiled from [33,35,50,51]. Discrepancies among references are shown in parentheses, suggesting partial modifications or modifications at certain conditions.

It has been shown in *V. cholerae* that hypomodification, s^4^U specifically, modulates tRNA abundance of a subset of tRNAs in the stationary phase. Interestingly, tRNA^Gln1^ is the only tRNA whose abundance is significantly reduced in the log phase [16]. We evaluated tRNA^Gln1^ and tRNA^Gln2^ abundance in *E. coli* WT, Δ*truA*, and Δ*mnmA* strains in both log and stationary phases. Bulk tRNA purified from samples harvested at 10, 20, 30, 45, and 60 minutes after addition of rifampicin that inhibits newly RNA transcription. tRNA abundance was detected through Northern blotting using probes for tRNA^Gln1^ and tRNA^Gln2^. We found no significant changes in tRNA abundance suggesting these tRNAs, tRNA^Gln1^ and tRNA^Gln2^, are stable in Δ*truA* and Δ*mnmA* (**Fig. S4**). However, we cannot rule out that these could be unstable when both genes are deleted but this cannot be tested as the combination of disruption is not viable.

### The absence of unique tRNA modifications does not drastically influence tRNA levels in *E. coli* K12

One possible cause for observing synthetic lethality phenotypes when combining tRNA modification genes is that the tRNAs lacking the two modifications are degraded by RNase as recently seen in *Vibrio* or *Salmonella* [16,36]. Using AQRNA-seq, a tRNA-seq method that can measure absolute levels [37], we quantified levels of all *E. coli* tRNAs in cells grown to mid-log phase (OD_600nm_ of ~0.5) in 10 of the 16 mutant strains that led to synthetic lethality phenotypes in certain combinations (**Fig. S5** and **Supplementary data S3**)

We found that the levels of 11 tRNA decreased significantly in tRNA modifications mutants (**Supplementary data S3F**), 9 of those were modified by the corresponding enzymes, *dusB*, *thiI*, *trmJ*, *miaA*, *mnmC*, *mnmG*, and *truA* (**Supplementary data S3F**) but none matched with the synthetic lethal combinations.

### Synthetic lethal screens lead to the discovery of condition-dependent synthetic lethality

As discussed above, deletion of *rlmN*, *mnmA*, or *trmJ* resulted in synthetic lethal combinations with several other tRNA modification mutants (**Fig. 2**). These four genes encode enzymes that modify a common substrate: tRNA^Gln1^_UUG_ (**Fig. 4A**) but did not lead to any growth defects when deleted in pairwise combinations in the initial screen performed on LB (**Fig. 2**). To study these strains further, we generated the 6 pairwise deletion strains through P1 transductions (**Table S1**). We tested the fitness of the double knockout pairs by determining the growth rate in LB at 37 °C (**Fig. S6A**). The Δ*rlmN*Δ*mnmA* and Δ*trmJ*Δ*mnmA* strains showed a slight growth defect, with Δ*rlmN*Δ*mnmA* exhibiting an increased lag phase compared to Δ*rlmN* and Δ*mnmA* strains (**Fig. S6A**). We then conducted a screen to determine whether growth of the Δ*rlmN*Δ*mnmA* strain could be affected under specific growth conditions by varying the carbon source (glucose, xylose, acetate, and glycerol) and the incubation temperature (30 °C, 37 °C, or 42 °C) (data not shown). We found that Δ*rlmN*Δ*mnmA* mutant showed a slight growth defect when cultured on acetate and xylose containing media at 37 °C (**Fig. S6B**). All strains carrying the *mnmA* deletion failed to grow on acetate at high temperature but only the Δ*rlmN*Δ*mnmA* strain failed to grow on xylose at 42 °C (**Fig. 5A-C**). Because tRNA^Gln1^ is the only tRNA modified with both s^2^U and m^2^A37 (**Fig. 4**), we tested whether its overexpression could suppress the temperature sensitivity (TS) of the Δ*rlmN* Δ*mnmA* strain when grown in M9 supplemented with with xylose (0.4%) as the carbon source. We found that neither tRNA^Gln1^_UUG_ nor tRNA^Gln2^_CUG_ suppressed the TS phenotype in xylose (**Fig. S7A**).

**Figure 5.**
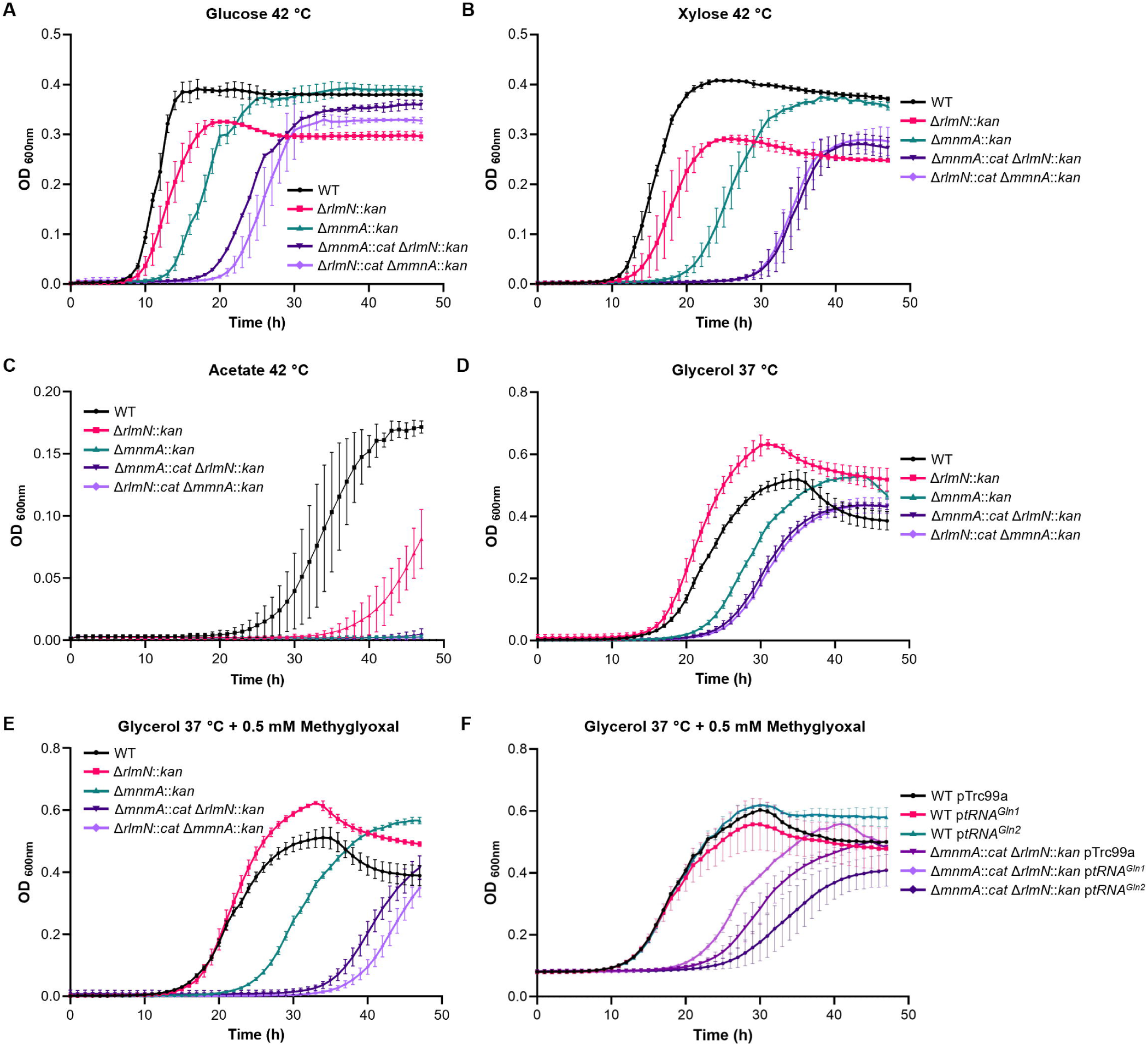
Growth phenotypes of the *rlmN mnmA* strain. Growth of the WT, *rlmN*, *mnmA* and *rlmN mnmA* strains at 42 °C in M9 supplemented with 0.4% glucose (**A**), 0.4% xylose (**B**), and 0.4% acetate (**C**) as carbon source. (**D-E**) Growth of the WT, *rlmN*, *mnmA,* and *rlmN mnmA* strains at 37 °C in M9 with glycerol (0.4%) as carbon source and with or without methylglyoxal (0.5 mM). (F) Growth of WT and *mnmA rlmN* strains expressing tRNA^Gln1^ (p*tRNA^Gln1^*) or tRNA^Gln2^ (p*tRNA^Gln2^*) in M9 with glycerol (0.4%) as carbon source, methylglyoxal (0.5 mM), IPTG (25 µM), and ampicillin (100µg/mL). The empty plasmid pTrc99a was used as a control. The average OD of three biological replicates were plotted with error bars representing the standard deviation.

It has been previously reported that excess xylose can lead to increased production of methylglyoxal [38]. The two methylglyoxal detoxification pathway genes *gloA* and *gloB* preferentially use the UUG codons decoded by tRNA^Gln1^ (**Supplementary data S4**). When grown in M9 supplemented with 0.4% glycerol and 0.5mM methyglyoxal, Δ*rlmN*Δ*mnmA* shows a growth defect compared to WT and the single knockouts (**Fig. 5D-E**). We found that over expression of tRNA^Gln1^_UUG_ moderately suppressed the phenotype in the double knockouts (**Fig. 5F**). However, we observed growth inhibition of tRNA^Gln2^ overexpression uniquely in the same strain (**Fig. S7B**).

Together, these results demonstrate that loss of multiple modifications affecting tRNA^Gln1^ did not invariably lead to growth defects under standard conditions but instead exhibited condition dependent phenotypes that became apparent under specific metabolic and environmental stresses. In particular, the Δ*rlmN* Δ*mnmA* strain displayed sensitivity to elevated temperature, alternative carbon sources, and methylglyoxal, phenotypes that could be partially modulated by altering tRNA dosage. These findings reveal that synthetic interactions among tRNA modification genes can manifest as stress specific growth defects rather than unconditional lethality.

## Discussion

By systematically constructing all pairwise combinations of 16 *E. coli* tRNA modifications genes, we identified specific phenotypes caused by the disruption of two tRNA modification genes (**Fig. 2)**. Twenty cases of negative epistasis were identified and 8 of those involved *truA*. Three of 5 synthetic lethal pairs (*truA*/*rlmN*, *rlmM*/*miaA* and *trmJ*/*truA*) had showed negative genetic interaction scores in the synthetic lethality screens that focuses on translation genes from the Emili laboratory, but these were under their high-confidence cut-off [22]. Also, several tRNA modification genes including *mnmA* were not present in their initial target list. Hence, our results reinforce the value of orthogonal approaches to measure essentiality as we use P1 transduction efficiency, while the Emili group study used colony size [22].

The roles of specific tRNA modifications seem to vary greatly with both the organism, and the growth conditions. We did not observe in *E. coli* the synthetic lethality of *thiI* with *miaA*, *truB* and *trmA* observed in *V. cholerae* [16]. We did see that the combination of *thiI* and *mnmG* deletions was detrimental (**Fig. 2**). Similarly, in Salmonella the absence of Ψ39-40 in a *truA* mutant leads to the accumulation of tRNA halves but this is not the case in *E. coli* as already reported by the Wolin laboratory [17] (**Supplementary data S3**). Growth conditions are also critical to reveal the physiological roles of tRNA modifications. For example, the degradation of tRNA^Tyr^ by the degradosome in the absence of ThiI in *V. cholerae* is mainly observed in stationary phase [16]. In the same organism the growth defects of the *thiI miaA*, *thiI truB* and *thiI trmA* strains is more pronounced at 37 °C compared to 25 °C [16]. In our study, the growth defects of the *rlmN mnmA* strain were dependent on both the carbon source and the temperature (**Fig. 5**). Hence to fully explore the physiological roles of tRNA modification will require the use of genome wide TnSeq screens using tRNA modification deletion strains as starting points as the technology to perform such screens has recently become available [39]. These whole genome screens could also identify interactions between tRNA modifications genes and other genes related or not to translation. For example, targeted studies have shown that deleting *truA* or *iscS* is synthetic lethal in combination with the Δ*rnhAB* deletion that is deficient in RnaseH revealing that translation efficiency affects DNA replication [20].

The presence of Ψ39-40 in the ASL dependent on TruA is emerging as key quality control marker for tRNA integrity in different organisms but possibly through different mechanisms. We also found that in *E. coli* K12 tRNA^Gln1^ seems to be particularly dependent on tRNA modifications for stability and further studies are required to identify the players involved even if it is highly probable that the degradosome is involved. Finally, this work reinforces the power of using epistatic interactions screens in general to discover novel molecular mechanisms even in a very well-studied organism like *E. coli* K12.

## Methods

### Bioinformatics

Information on *E. coli* K12 genes was extracted from EcoCyc [40] (NZ_CP009273) and the resources of NCBI including PubMed [41]. Codon usage was performed using the Codon UTilization tool (CUT) tool [42]. Molecular graphics and analyses performed with UCSF ChimeraX [43].

### Bacterial strains, plasmids and media

Luria-Bertani (LB) broth and agar (tryptone 10 g/L, yeast extract 5 g/L, sodium chloride 10 g/L, Fisher Scientific BP1426-2 and BP1425-2) were routinely used for growth of *E. coli* cells at 37 °C from frozen stocks. M9 minimal medium (12.8 g/L Na_2_HPO_4_·7H_2_O, 3.0 g/L KH_2_PO_4_, 0.5 g/L NaCl, 1.0 g/L NH_4_Cl, 2.0 mM MgSO_4_, 0.1 mM CaCl_2_) was used with 0.4% glucose, xylose, acetate, glycerol, or succinate as carbon sources. When required, 50 µg/mL of kanamycin (KAN), 200 µg/mL Ampicillin (AMP), 20 µg/mL of chloramphenicol (CHL), 2.5 µM-1mM Isopropyl β-D-1-thiogalactopyranoside (IPTG), methylglyoxal (0.5 mM) or arabinose (0.02%) were added to the medium. Bacterial growth was followed in a Bioscreen C or Bioscreen G automated growth curve analyzer (Growth Curve, USA) by measuring OD_600nm_ over a period of 18-24 hours.

The bacterial strains used in this study are listed **Table S1**. Most are from the Keio collection [44] and from a set of Hfr strains containing a single deletion of tRNA modifying genes replaced by cat cassette (H. Mori) [45]. Markerless deletions strains were generated by FLP mediated excision of the kan cassette through pCP20 [46] for a subset of Keio collections strains. Plasmid transformations were performed by electroporation using manufacturer’s settings (Bio-Rad Micropulser). All strains carrying two tRNA modification alleles were constructed by P1 transduction as described below. All the strains were validated for the correct genotype using the primers listed on **Table S2.**

The fragments encoding tRNA^Gln1^ and tRNA^Gln2^ were subcloned from pSADgln1,2 [47] into the EcoRI and PstI sites in pTrc99a yielding pJMB23 (pTrc99a-tRNA^Gln1^) and pJMB24 (pTrc99a-tRNA^Gln2^). pTrc99A, pJMB23, and pJMB24 were transformed into wildtype, Δ*mnmA::cat*Δ*rlmN::kan*, Δ*rlmN::cat*Δ*mnmA::kan*, and Δ*truA::kan* through electroporation. *truA* was amplified by PCR using *E. coli* BW25113 genomic DNA as template using a primer pair bearing 5′-EcoRI and 3′-SalI sites (**Table S2**). The PCR product was digested with EcoRI and SalI (NEB) and ligated (NEB T4 DNA ligase) into the corresponding sites in pBAD24 yielding pBAD-*truA*. The empty pBAD24 and pBAD-*truA* were transformed into wildtype and Δ*truA::kan* through electroporation.

### Synthetic lethality analysis by P1 Phage Transduction

P1vir phage [48] donor lysates were prepared from single deletion strains containing a chloramphenicol resistance cassette (cat) in place of the deleted gene [45]. The single deletion strains from the Keio collection [44] containing a kanamycin resistance cassette (kan) served as the recipient. Donor Lysates were prepared as previously described [49]. Briefly, donor strains were grown in P1-LB (LB supplemented with 5 mM CaCl_2_, 0.1% glucose, and 12.5mM of MgCl_2_) to an OD_600nm_ of 0.6, to which 40 µL of P1vir phage was added. Infection was allowed to proceed for 4 hours until cell debris was visible. The mixture was then centrifuged for 15 minutes at 4,000 rpm to separate the cell debris. The supernatant was passed through a 0.2 µM filter to remove any trace cell debris and residual bacteria. Chloroform (200 µL) was added to prevent bacterial growth in the lysate. Recipient strains were grown in P1-LB to an OD_600nm_ of 0.6. One mL was transferred into a 2 mL centrifuge tube and centrifuged at 13,000 rpm for 1 min. Cell pellets were resuspended in 100 µL P1-LB, to which 20 µL of the donor lysate was added. The transfection was allowed to proceed for 15 minutes with constant orbital shaking at 180 rpm and the reaction was terminated by adding 200 µL of 1M sodium citrate pH 5.5- and 1-mL LB. Transductants were allowed to recover for 2 h in 37 °C with shaking. Reaction was centrifuged at 13,000 rpm and the pellet was resuspended in 150 µL LB-citrate (LB supplemented with 100 mM sodium citrate pH 5.5) and plated on LB-agar plates supplemented with 50 μg/mL kanamycin and 25 μg/mL chloramphenicol for selection. Plates were incubated at 37 °C for 16-20 h. The transduction efficiency was assessed by quantifying colony-forming units (CFU) comparing to a wildtype (WT) recipient. We used the following scale to categorize transduction efficiencies: 1-10 colonies – defective, zero colonies – lethal, small colonies. A second round of P1 transductions was performed independently to verify the 29 pairs categorized as synthetic lethal pairs from the initial round (**Supplementary Data S2**).

### tRNA purification

*E. coli* cultures (10 mL) grown to OD_600nm_ ~0.6 were harvested by centrifugation 4 °C for 10 min at 3700 xg. Pellets were resuspended in 1 mL of Trizol (Life Technologies) and agitated on a rocking shaker for 2-4 h. Small RNA was extracted using a Purelink miRNA Isolation kit (Invitrogen) and RNA eluted in 50 μL of RNase-free water. Following RNA quantification by Nanodrop, samples with A_260_/A_230_ ratios <1.6 were cleaned using an RNA Clean and Concentrator Kit (Norgen Biotek).

### tRNA stability assay

*E. coli* cultures (Δ*mnmA*::*kan* (JTB1119), Δ*truA*::*kan* (JW2315), and WT BW25113 (JTB1001)) were grown to log phase (OD_600nm_ = 0.5-0.6) and stationary phase (OD_600nm_ = 0.8-0.9). Cultures were spiked with 100 µg/mL final concentration rifampicin. Two mL samples were harvested at specific time points after rifampicin spike (10, 20, 30 and 45 min for log and stationary phases, and 60 min for the stationary phase only) for bulk tRNA extraction and purification. tRNA was purified using PureLink miRNA Isolation Kit (Invitrogen). tRNA degradation was quantified through Northern blotting using probes specific to the TΨC-loop of tRNA Gln1 and tRNA Gln 2. 5S rRNA was used as loading reference. One microgram of purified tRNA, alongside Pre-stained Marker of Small RNA plus (BDL DM253), was run on a Mini-PROTEAN 10% TBE-Urea Precast Gel (Bio-Rad) at 120 V for 2.5 h in an ice bath with constant stirring. Gel bands were transferred to a Biodyne B Modified Nylon Membrane (Thermo-Scientific) using a Bio-Rad Trans-Blot SD Semi-Dry Transfer Cell, at 10 V for 15 min, followed by UV Crosslinking using UV Crosslinker FB-UVXL-1000 (Fisher Scientific) for 1200 μs at optimal settings. The membrane was cut horizontally at the 75 bp marker. The upper half was probed for 5s rRNA while the lower half was probed for either tRNA^Gln1^ or tRNA^Gln2^. Hybridization was run overnight at 42 °C with constant rocking. Membranes were developed using the Chemiluminescent Nucleic Acid Detection Module Kit (Thermo Fisher Scientific) and visualized using iBright FL1500 Imaging System (Thermo Fisher Scientific).

### Mass Spectrometry Analysis of tRNA

RNA in each sample (1.8 µg) was hydrolyzed in a 30 µL digestion reaction containing 2.49 U benzonase, 3 U CIAP (calf intestinal alkaline phosphatase), 0.07 U PDE I (phosphodiesterase I), 0.1 mM deferoxamine (antioxidant), 0.1 mM butylated hydroxytoluene (antioxidant), 3 ng coformycin (adenosine deaminase inhibitor), 25 nM [^15^N_5_]-2’-deoxyadenosine (internal standard), 2.5 mM MgCl_2_ and 5 mM Tris-HCl buffer pH 8.0. The digestion mixture was incubated at 37 °C for 6 h. After digestion, all samples were loaded into sample vials containing an Agilent sample vial insert and analyzed by chromatography-coupled triple-quadrupole mass spectrometry (LC-MS/MS). For each duplicate sample, 600 ng of hydrolysate was injected twice as technical replicates on a Waters Acuity BEH C18 column (50 × 2.1 mm inner diameter, 1.7 µm particle size) eluted with the following buffer gradient of Buffer A (0.02% formic acid) and Buffer B (0.02% formic acid in 70% acetonitrile) was used for elution of ribonucleosides at 0.3 mL/min according the time frame given **Table S3**. The HPLC column was coupled to an Agilent 1290 HPLC system and an Agilent 6495 Triple Quadrupole LC/MS spectrometer and eluted at a flow rate of 0.3 mL/min and 25 °C, with an electrospray ionization source in positive mode with the following parameters: drying gas temperature, 200 °C; gas flow, 11 L/min; nebulizer, 20 psi; sheath gas temperature, 300 °C; sheath gas flow, 12 L/min; capillary voltage, 3000 V; nozzle voltage, 0 V. The first and third quadrupoles (Q1 and Q3) were fixed to unit resolution, and the modifications were quantified by pre-determined molecular transitions. The dwell time was 500 ms, the fragmentor voltage was 166 V, and the cell accelerator voltage was 5 V for each nucleoside. Multiple reaction monitoring (MRM) mode was used for detecting product ions derived from precursor ions for the ribonucleosides, with instrument parameters including the collision energy (CE) optimized for maximal sensitivity for the modification. Synthetic standards (except for oQ) were used to define HPLC retention times of RNA modifications. The retention time, *m/z* of the transmitted parent ion, and *m/z* of the monitored product ion for each ribonucleoside are detailed in **Table S4**. All MRM peak integrations were manually checked.

### Determination of absolute tRNA levels by AQRNA-seq

#### AQRNA-seq library preparation

AQRNA-seq analysis of tRNA isoacceptors was performed as described in detail elsewhere [37]. Briefly, purified small RNA (50 ng) was mixed with an 80-mer spike-in RNA oligonucleotide internal standard (DNA and RNA oligonucleotide sequences in **Table S5**). The mixture was dephosphorylated shrimp alkaline phosphatase (rSAP, NEB) in NEB T4 RNA ligase buffer at 37 °C for 30 min, followed by heat inactivation at 65 °C for 5 min and cooling on ice. Linker 1 ligation was performed by adding Linker 1 (100 pmol/ul; **Table S5**), ATP (10 mM, NEB), T4 RNA ligase buffer (NEB), T4 RNA ligase 1 (30U/ul, NEB), water and PEG8000 (NEB) to the reaction mixture. Following incubation at 25 °C for 2 h and then at 16 °C overnight, the ligation product was purified using the Zymo Oligo Clean & Concentrator kit (Zymo Research, D4060). The sample was eluted in water and kept on ice prior to Bioanalyzer analysis (Agilent, small RNA kit) or used directly in the demethylation step. Next, the DNA adenylated oligonucleotide adenylate intermediate was de-adenylated (RNA sample, NEB Buffer 2, and 5’-deadenylase (NEB); incubation at 30 °C for 1 h) and removed together with unused Linker 1 in a 30 min incubation with exonuclease RecJ (NEB). The reaction was stopped by heating at 65 °C for 20 min and products purified using a DyEx spin column (Qiagen). Reverse transcription was performed by mixing purified RNA with RT-primer (2 pmol/ μl) and dNTPs (10 mM each) and heating to 80 °C for 2 min, followed by cooling on ice. PrimeScript Buffer (Clontech), RNase Inhibitor (NEB), and PrimeScript Reverse Transcriptase (Clontech) were added with incubation at 50 °C for 2 h followed by enzyme inactivation at 70 °C for 15 min. The RNA strand was removed by adding NaOH (5 M) with incubation at 90 °C for 3 min, followed by neutralization with HCl (5M) and cDNA purification using the Zymo Oligo Clean & Concentrator kit (Zymo Research). The sample was eluted with water and vacuum concentrated to 5 ul. The purified cDNA was ligated to Linker 2 (**Table S5**) with T4 DNA Ligase Buffer (NEB), ATP (NEB), T4 DNA ligase (NEB), and PEG8000 (NEB) at 16 °C overnight. Ligated product was purified using the Zymo Oligo Clean & Concentrator kit and eluted in 16 μl of water. Excess Linker 2 and adenylated linker products were removed by RecJ and the reaction was stopped through heat inactivation at 65 °C for 20 min. Purified cDNA was then amplified by PCR in seqAMP DNA polymerase buffer (Clontech) with PCR primer F and R with unique sequencing barcodes (**Table S5**), seqAMP DNA polymerase (Clontech), and water. PCR was performed with an annealing temperature of 58 °C and 13 reaction cycles followed by agarose gel resolution and extraction using a standard gel purification kit (QIAquick Gel Extraction Kit, Qiagen). The gel-extracted samples were mixed for multiplexing and submitted for Illumina NextSeq sequencing.

#### AQRNA-seq data processing

Illumina sequencing reads were assessed using a quality control pipeline and the data subsequently processed using a workflow described in detail elsewhere [37]. Briefly, adapter sequences were removed from forward and reverse reads using fastxtoolkit (v0.013). Sequences were blasted against a reference library (including the 80-mer internal standard) using blast (v2.6.0) with the parameters: blast -perc identity 90 -word_size 9 -dust no -soft_masking false. Sequences corresponding to duplicate tRNA genes and tRNA pseudogenes were removed to eliminate redundant entries and reduce the incidence of ambiguous or false positive matches. The terminal (3’) CCA sequence was added to tRNA sequences where it is not genomically encoded. For each tRNA and control sample, forward and reverse reads were merged by integrating their start and end positions to generate new start and end positions that reflect their combined coverage. Using the python script cull.py, multiple alignments were reduced by ranking all the alignments for a given read by their e-value and retaining only the alignment with the lowest e-value, retaining forward and reverse reads that matched the same target. Uniquely mapped reads were then tabulated and counted.

## Acknowledgments

We thank Thomas Begley (University at Albany) for running the CUT analysis on the *E. coli* genome and William McClain for sharing the pSADgln1,2 plasmid.

## Data availability Statement

All raw data have been deposited in public repositories: LC-MS/MS RNA data have been deposited to the ProteomeXchange Consortium via the PRIDE [52] partner repository with the dataset identifier PXD078891; AQRNA-seq data have been deposited NCBI’s Gene Expression Omnibus [53] and are accessible through GEO Series accession number GSE331519 (https://www.ncbi.nlm.nih.gov/geo/query/acc.cgi?acc=GSE331519).

## Statement of AI use

Microsoft 365 Copilot was used to assist with language editing and clarity. The authors take full responsibility for the content.

